# Using Human Induced Pluripotent Stem Cell Derived Organoids to Identify New Pathologies in Patients with PDX1 Mutations

**DOI:** 10.1101/2021.04.25.441341

**Authors:** Mansa Krishnamurthy, Daniel O Kechele, Taylor Broda, Xinghao Zhang, Jacob R Enriquez, Heather A McCauley, J Guillermo Sanchez, Joseph Palermo, Margaret Collins, Inas H Thomas, Haley C Neef, Amer Heider, Andrew Dauber, James M Wells

## Abstract

Two patients with mutations in *PDX1* presented with pancreatic agenesis, chronic diarrhea and poor weight gain, the causes of which were not identified through routine clinical testing. We generated patient derived organoids as a novel diagnostic strategy and observed that *PDX1*^188delC/188delC^ antral organoids convert to an intestinal phenotype, while intestinal organoids undergo gastric metaplasia with significant reduction in enteroendocrine cells. This prompted a re-examination of gastric and intestinal biopsies from both *PDX1*^188delC/188delC^ patients, which recapitulated organoid phenotypes. Antral biopsies had increased parietal cells and lacked G-cells suggesting loss of antral identity. These patients will now be monitored for the progression of metaplasia. This study demonstrates the utility of organoids for patient diagnoses and treatment.

## Introduction

Pancreatic and duodenal homeobox gene-1 *(PDX1)* is a parahox transcription factor that is essential for embryonic development of the pancreas in mice and humans (1–6). Patients bearing homozygous loss-of-function mutations in *PDX1* are born with pancreatic agenesis, neonatal diabetes and gallbladder agenesis (7,8). Additional phenotypes have been identified in *Pdx1* null mice, including a contorted pylorus, ectopic biliary duct epithelium in distal regions of the duodenum and a reduction in some enteroendocrine cells (EECs) (5,6,9). However, none of these pathologies have been identified in humans, possibly due to limitations of available diagnostic tests. The disruptions in boundary formation within gastrointestinal (GI) epithelium certainly pose a potential risk for the development of GI cancer in this patient population. Moreover, abnormalities in EEC development and function can negatively affect hunger, satiety, digestion and absorption. Thus, it is essential to identify the exact effects of *PDX1* mutations in humans to provide optimal care for these patients.

Over the past decade, we have developed several human organoid models that have allowed for unprecedented modeling of human GI development and function (11–15). Organoids are three-dimensional human tissues that function in a manner similar to native organs and thus allow for unparalleled mechanistic studies of patient pathophysiology in the laboratory. For example, organoid model systems bioengineered from patient derived induced pluripotent stem cells (iPSCs) with CRISPR/Cas9 technology for isogenic correction of mutations affords us a unique opportunity to diagnose new pathologies, mechanistically interrogate human diseases, and individualize patient therapies.

We have identified two patients with a homozygous frameshift mutation in *PDX1*, causing complete loss of functional protein. In addition to having diabetes and exocrine pancreatic insufficiency, these patients had additional GI symptoms including chronic diarrhea and poor weight gain. Initial endoscopies and biopsies failed to identify GI pathologies. We took a novel diagnostic approach and used patient derived iPSCs to generate gastric and intestinal organoids to diagnose how mutations in *PDX1* cause a myriad of GI pathologies. This *in vitro* modeling led us to re-examine the patients’ clinical samples and confirm the *in vivo* pathophysiology thus impacting their clinical care. For the first time, we demonstrate that lack of PDX1 in humans results in gastric and intestinal metaplasia and increased inflammation in the GI tract. We also show that loss of PDX1 decreases key EECs including somatostatin, PYY, Ghrelin and GIP and an overall loss of antral identity. This study represents a paradigm shift in how we use human organoids to identify and model complex pathologies in patients and alter patient care.

## Methods

### Patient Phenotypes

Patient A: A male infant with a confirmed homozygous mutation in *PDX1* (c.188delC (p.P63fs)) was identified via clinical testing. The study protocol conformed to the ethical guidelines of the 1975 Declaration of Helsinki and was approved by the Institutional Review Board of Cincinnati Children’s Hospital Medical Center. Written informed consent was obtained from the subject’s legal guardian. He was born small for gestational age with severe intra-uterine growth restriction (birth weight 1125 grams (−3.8SD), birth length 36.2 cm (−4.48SD), head circumference 36.2 cm (−3.13 SD)). At birth, he was noted to be hyperglycemic with blood glucose measurements >600 mg/dL. He was diagnosed with neonatal diabetes and initiated on insulin therapy. Additional GI work up demonstrated partial gallbladder agenesis and pancreatic agenesis causing exocrine pancreatic insufficiency. Pancreatic enzyme replacement therapy was started immediately, and a G tube was placed to ensure adequate caloric intake in the setting of feeding difficulties. At two years of age, the G tube was removed. Esophagogastroduedonsocpy (EGD) was performed at 24 and 40 months of age with routine biopsies obtained from the antrum and duodenum. The EGD at 24 months to close his gastrocutaneous fistula found mucosal nodularity in the antrum and pre-pyloric region. The duodenal mucosa was normal. Luminal narrowing was encountered at the D1/D2 transition. At 40 months of age, a repeat EGD found normal appearing mucosa in the stomach and duodenum without duodenal narrowing. He is currently managed on an insulin pump, pancreatic enzymes and proton pump inhibitor therapy.

Patient B: Similarly, another male infant with a confirmed homozygous mutation in *PDX1* (c.188delC(p.P63fs)) was born at 37 weeks with severe intra-uterine growth restriction (birth weight 1560 g (−3.35 SD)) with pancreatic agenesis and gallbladder agenesis. This clinical presentation has been previously described (9). He is managed on insulin injections, pancreatic enzymes, proton pump inhibitor, cyproheptadine, and lactobacillus rhamnosus.

### Pluripotent stem cell culture and directed differentiation into pancreatic cells, HAGOs and HIOs

iPSC lines were generated from Patient A as previously described (10). iPSC clones were maintained on hESC-qualified Matrigel (Corning) in mTeSR1 media (STEMCELL Technologies) and confirmed to be pluripotent, karyotypic normal and mycoplasma negative.

Human antral organoids, intestinal organoids, and pancreatic endocrine cells were generated as previously reported (10–14). For intestinal organoid generation, the AggreWell 400 (StemCell Technologies) system was used to generate spheroids. Plates were rinsed with anti-adherence rinsing solution. Cells were treated with Accutase (StemCell Technologies), rinsed in gut media and plated at a density of 3.6 million cells per well. The following day, spheroids were re suspended in Matrigel and were subject to intestinal protocol as described (10). Intestinal organoids were transplanted under the kidney capsule of immune deficient NOD.Cg-*Prkdc^scid^Il2rg^tm1Wjl^*/SzJ (NSG) mice and allowed to mature for 10 weeks as previously described (13).

### CRISPR Correction

CRISPR-Cas9 was used to correct a mutation in the *PDX1*^188delC/188delC^ iPSC line. The guide RNA (CTCGTACGGGGAGATGTCCG) targeting the mutation site was designed according to the web tool (http://CRISPOR.org) (15). The complementary DNA oligos were cloned as described (16). A phosphorothioated ssDNA donor oligo (GCCCTGGGCGCGCTGGAGCAGGGCAGCCCtC<u>C</u>GGAtATCTCCCCGTACGAGGTGCCtCCaCtCGCCGACGACCCCGCGGTGGCG) was designed to include the nucleotide insertion (underlined) according to published methods (17). iPSCs were reverse transfected with plasmid and donor oligo using TranIT-LT1 (Mirus). GFP-positive cells were isolated by FACS and replated at in Matrigel (Corning). Single clones were manually excised for genotyping, expansion, and cryopreservation. Correctly targeted clones were identified by PCR, enzyme digestion and Sanger sequencing.

### qPCR

RNA was extracted using the Nucleospin RNA extraction kit (Macharey- Nagel) and reverse transcribed into cDNA as previously described (18). Primer sequences are listed in Supplemental Table 2. qPCR was performed as previously described (18). Relative expression was determined using the ΔΔCt method and normalizing to PPIA (cyclophilin A). Organoid samples from at least three independent passages were used for quantification.

### Immunofluorescence Staining

Immunofluorescence staining on paraffin embedded tissues was performed as previously described (12). Primary and secondary antibodies are listed in Supplemental Table 1. Images were acquired using a Nikon A1 GaAsP LUNV inverted confocal microscope and were analyzed using NIS Elements (Nikon). Organoid samples from at least three independent passages were used for quantification. Data were quantified as number of positive cells divided by the entire epithelium labelled by either CLDN18, CDH17, Beta-Catenin or CDH1.

### Statistics

Data is presented as the mean +/- SEM. Significance was determined using appropriate tests in GraphPad Prism version 8, with *p<0.05, **p<0.01, ***p<0.001.

## Results

### Loss of PDX1 results in gastritis, intestinal and gastric metaplasia in the antrum and duodenum

Given that *PDX1^1^*^88delC/188delC^ patients had unexplained chronic diarrhea and weight loss, we developed an organoid-based strategy to perform deep diagnostic analysis of the stomach and intestine. As a control we confirmed that iPSC lines from a *PDX1^1^*^88delC/188delC^ patient was unable to generate pancreatic tissue consistent with pancreatic agenesis (Sup Fig 1). We generated antral and intestinal organoids using iPSC lines from the *PDX1*^188delC/188delC^ patient and an isogenic iPSC line where we corrected the *PDX1* point mutation using CRISPR/Cas9, thus restoring expression of PDX1 protein. While no macroscopic differences were observed between control and *PDX1*^188delC/188delC^ antral and intestinal organoids (data not shown), the *PDX1*^188delC/188delC^ organoids demonstrated broad dysplasia, with stomach tissue appearing in the intestinal organoids and vice versa (Figure 1). Intestinal markers upregulated in the antral organoids of the *PDX1^1^*^88delC/188delC^ patient included Cadherin 17 (CDH17), Villin1 (VIL1), CDX2, and MUC2 and decreased gastric markers Claudin 18 (CLDN18) and MUC5AC (Figure 1A,C, Sup Fig 1, 2). Conversely, gastric markers present in the intestinal organoids of *PDX1*^188delC/188delC^ patients included CLDN18 and MUC5AC. Moreover, regions of metaplasia in antral and intestinal organoids demonstrated a complete conversion from one epithelial type to the other (Figure 1B, D, Sup Fig 1, 2). These phenotypes were rescued in *PDX1* CRISPR corrected organoids indicating that the PDX1 mutation is unambiguously causing the metaplastic phenotypes. Together these findings suggest that loss of PDX1 results in gastric and intestinal metaplasia in humans.

**Figure 1:**
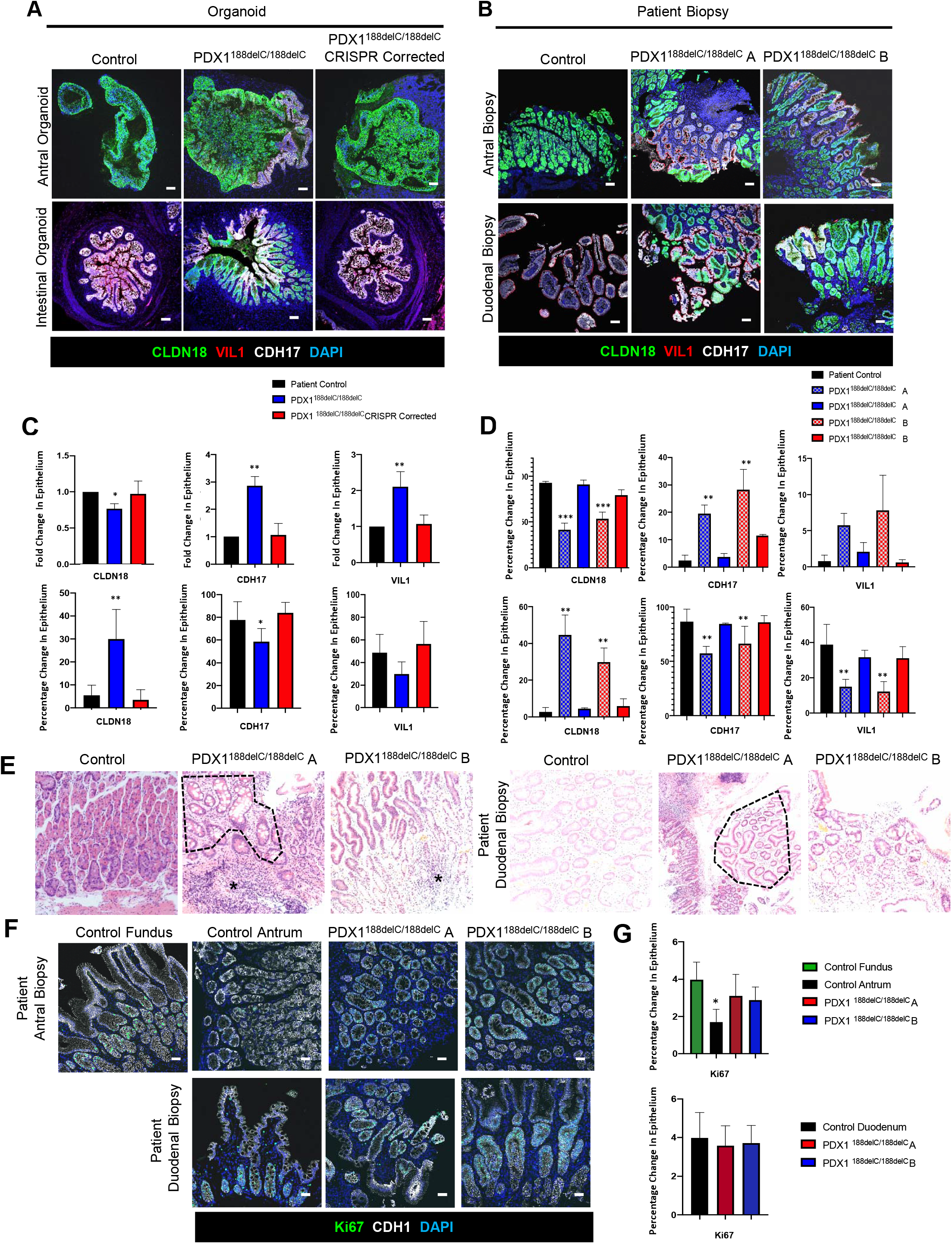
Loss of PDX1 results in intestinal or gastric metaplasia in the stomach and intestine, respectively. **(A-B)** Immunofluorescence and **(C-D)** quantification of staining in control, PDX1^188delC/188delC^, and PDX1^188delC/188delC^ CRISPR Corrected **(A)** antral and post-transplantation intestinal organoids and (**B**) antral and duodenal biopsies from control, PDX1^188delC/188delC^ A and PDX1^188delC/188delC^ B biopsies stained with gastric epithelial marker CLDN18 and intestinal epithelial markers CDH17 and VIL1. **(E)** Histological staining of antral and duodenal biopsies with regions of metaplasia encircled and inflammatory foci marked with asterisk. **(F)** Immunofluorescence and **(G)** quantification of control fundus, antrum and duodenum, and PDX1^188delC/188delC^ A and PDX1^188delC/188delC^ B patient antral and duodenal biopsies stained with proliferation marker Ki67 and epithelial marker CDH1. All sections counterstained with nuclear DAPI. Scale bars represent 100 μm.

Based on the above organoid findings, antral and intestinal biopsy samples from both *PDX1*^188delC/188delC^ patients were re-examined to look for evidence of metaplasia. Not only did biopsies from both patients demonstrate gastric and intestinal metaplasia, (Figure 1E-H, Sup Fig 1, 2), biopsies from both *PDX1*^188delC/188delC^ patients revealed overt inflammation in the antrum (Figure 1I-J) especially in regions close to intestinal metaplasia. Additionally, the metaplastic areas in the antrum had increased KI67 proliferation when compared to control patients (Figure 1K-N) suggesting these areas are mitotically active. These findings indicate that these patients are at increased risk for developing cancer. Thus the clinical team caring for these patients now provide additional screening to monitor for progression of inflammation, intestinal and gastric metaplasia.

### Loss of PDX1 results in loss of Antral identity and expression of fundic markers

PDX1 is normally expressed in the antrum and not in the fundus of the stomach. In the *PDX1*^188delC/188delC^ antral organoids and patient biopsies, loss of PDX1 expression is linked to loss of antral markers such as gastrin. Instead, antral organoids and biopsies are more fundic in nature, with an increase in parietal cells (ATP4B), Ghrelin positive cells, and Chief cells expressing fundic specific pepsinogens (PGA3) (Figure 2 A-H, M). The aforementioned markers were expressed at levels similar to control fundus, indicating that loss of PDX1 results in either a failure of the antrum to form or a possible antral to fundic conversion. The loss of a functional antrum would be expected to have clinical consequences in these patients which include delayed gastric emptying and deranged parietal cell function due to loss of gastrin-producing cells. Moreover, we observed ATP4B+ parietal cells in the intestinal organoids and biopsies of *PDX1*^188delC/188delC^ patients (Figure 2 I-L) suggesting the metaplastic tissue was fundic in nature. The increase in parietal cells in the antrum and duodenum may also explain the chronic diarrhea and lack of weight gain noted in these patients. All organoid results were discussed in real time with the team of physicians managing these challenging patients. Patients were placed on proton pump inhibitors and are monitored for the development of new metaplasia. Given the importance of the antrum in gastric emptying and the regulation of gastrin on parietal cell function we will monitor patients for additional complications in moving forward.

**Figure 2:**
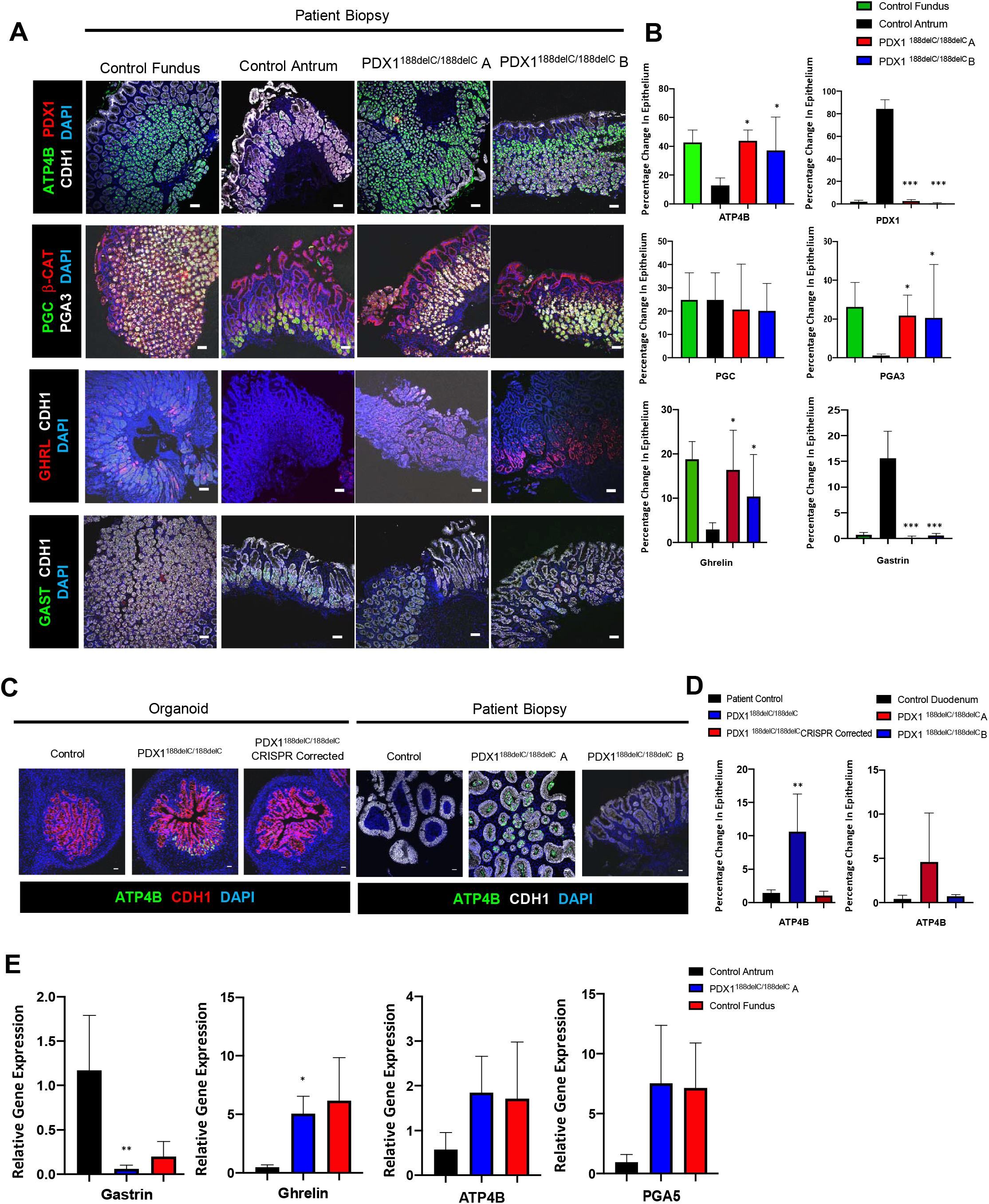
Loss of PDX1 results in loss of antral identity in the stomach. **(A)** Immunofluorescence and **(B)** quantification of staining in control fundic, antral, PDX1^188delC/188delC^ A and PDX1^188delC/188delC^ B biopsies stained with parietal cell marker ATP4B, transcription factor PDX1, pepsinogens PGA3 and PGC and hormones GAST and GHRL. **(C)** Immunofluorescence and **(D)** quantification of staining in control, PDX1^188delC/188delC^, PDX1^188delC/188delC^ CRISPR corrected post-transplantation intestinal organoids and control, PDX1^188delC/188delC^ A and PDX1^188delC/188delC^ B patient duodenal biopsies with gastric marker ATP4B and epithelial marker CDH1. All sections counterstained with nuclear DAPI. Scale bars represent 100 μm. (**E**) qPCR analyses of control antral and PDX1^188delC/188delC^ antral and control fundic organoids for GAST, GHRL, ATP4B and PGA5.

### PDX1 is required for normal enteroendocrine cell development in the Duodenum

EECs are nutrient sensing cells and in the duodenum are essential for nutrient absorption, motility, glucose homeostasis and satiety. Given its role in pancreatic endocrine cell development we surveyed EECs in the duodenum of *PDX1*^188delC/188delC^ organoids and biopsies. We note a significant reduction in EECs as marked by chromograninA (CHGA), as well as loss or reduced numbers of cells expressing the hormones SST, PYY, Ghrelin and GIP (Figure 3A-G) in both organoids and biopsies. We did not observe any changes in serotonin or gastrin in intestinal organoids or biopsies from our *PDX1*^188delC/188delC^ patient (Figure 3D-G) demonstrating that PDX1 is required for some, but not all EECs. This reduction may have direct clinical consequences on nutrient homeostasis including digestion and macronutrient absorption and may be contributing to the poor weight gain and persistent diarrhea observed in PDX1^188delC/188delC^ patients.

**Figure 3:**
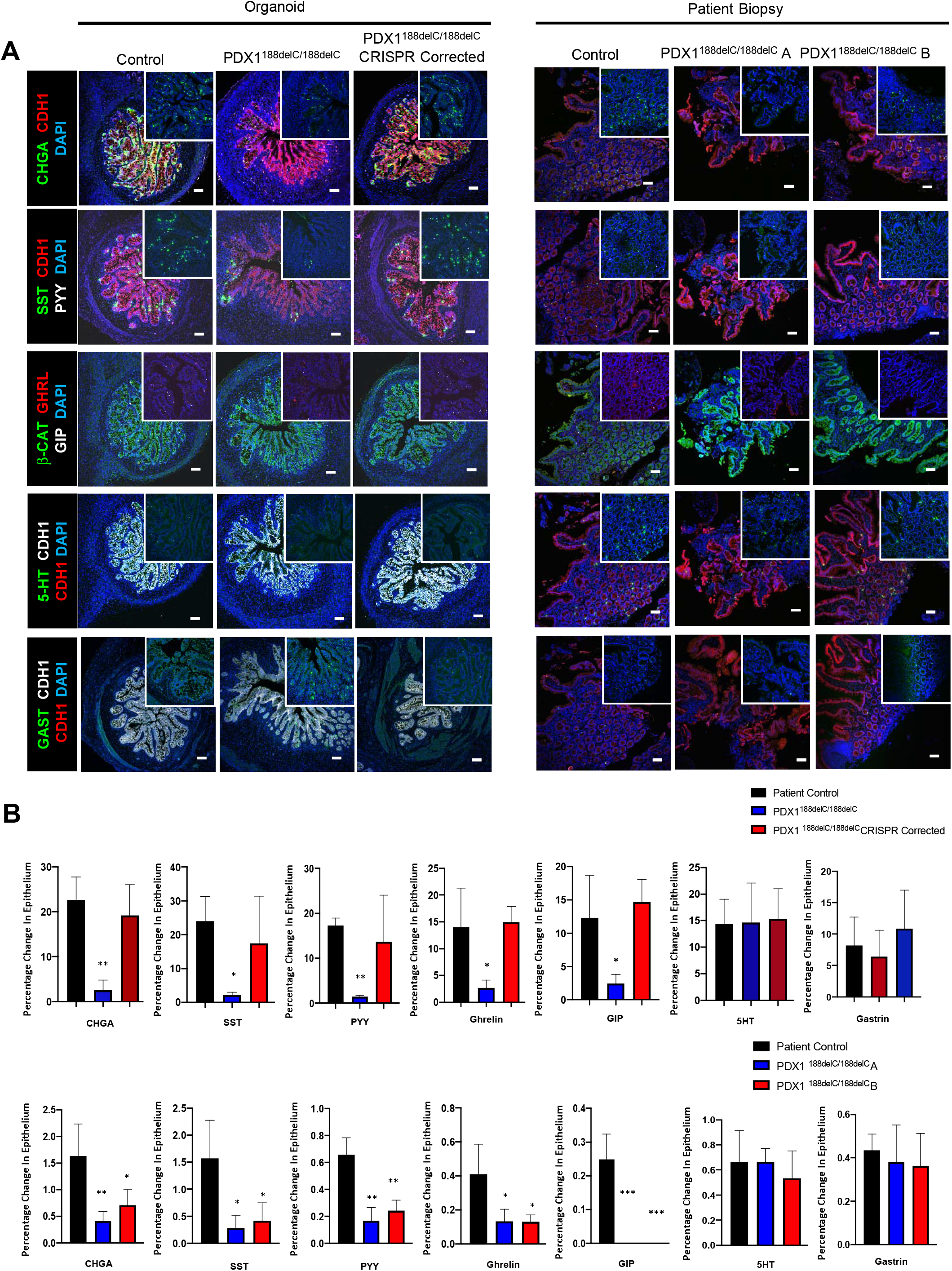
Loss of PDX1 decreases enteroendocrine cells in the intestine. **(A)** Immunofluorescence and **(B)** quantification of control, PDX1^188delC/188delC^, and PDX1^188delC/188delC^ CRISPR corrected post-transplantation intestinal organoids and control, PDX1^188delC/188delC^ A and PDX1^188delC/188delC^ B duodenal patient biopsies stained for secretory protein CHGA and enteroendocrine hormones SST, PYY, GHRL, GIP, 5HT and GAST. Insets show enteroendocrine cell staining at 20x magnification. All sections counterstained with nuclear DAPI. Scale bars represent 100 μm.

## Discussion

In this study, we generated patient derived GI organoids to uncover new pathologies in *PDX1*^188delC/188delC^ patients. We show that lack of PDX1 in humans results in an overall loss of antral identity, gastric and intestinal metaplasia and increased inflammation in the GI tract. We also show that PDX1 regulates expression of key EECs including somatostatin, PYY, Ghrelin and GIP in humans. Due to our discoveries using an organoid-based diagnostic strategy, these patients will now be monitored for the development of new areas of metaplasia and inflammation, that together increase the risk of development of gastric cancer. This study represents a paradigm shift in how we use human organoids to identify and model complex pathologies in patients and alter patient care.

Genetic ablation of *Pdx1* in mice results in loss of G cells in the antrum (3,4), which we also observed in *PDX1*^188delC/188delC^ gastric organoids and patient biopsies. However, *PDX1*^188delC/188delC^ patients still express gastrin in the intestine at similar levels as control, suggesting that the role of PDX1 for endocrine cell formation in the stomach is different from the intestine. Mice that lack gastrin have chronic gastritis, intestinal metaplasia of the stomach and eventually develop adenocarcinoma (19). *PDX1*^188delC/188delC^ patients also exhibit similar pathologies, suggesting that the absence of gastrin in the antrum may contribute to these pathologies. Given that gastrin secretion follows a meal and controls acid secretion, loss of gastrin and an increase in parietal cells might cause deranged gastric pH, improper digestion and bacterial overgrowth in the stomach and duodenum. We also noted the presence of parietal cells in the intestine in one of the *PDX1*^188delC/188delC^ patients which can explain the abdominal pain that these patients are experiencing and the gastritis noted in their biopsies. In patients who eventually develop gastric cancer, the extent of gastric intestinal metaplasia has been shown to be an important risk factor (20). Thus, from our studies, *PDX1*^188delC/188delC^ patients will be regularly monitored for further progression of metaplasia and inflammation as well as functional changes in the GI tract.

PDX1 is expressed only in the antrum of the stomach and not in the fundus (3,4). Interestingly, the *PDX1*^188delC/188delC^ patients demonstrate fundic marker expression in the antrum including increased Ghrelin, parietal and fundic pepsinogen cell populations, suggesting that PDX1 is essential for antral formation. Moreover, patients with Menetrier’s disease, have ectopic expression of PDX1 in the fundus, resulting in antralization and eventual expression of gastrin in fundic regions (21). Taken together with the findings of our study, these data suggest that PDX1 is required for antral formation during development. Interestingly, both *PDX1^188delC/188delC^* patients and those with Menetrier’s disease have diarrhea and difficulty with weight gain. This indicates that developmental disruptions in stomach formation may have significant downstream consequences including improper digestion and gastric emptying.

In summary, we have used a novel diagnostic approach with patient derived human GI organoids to uncover a myriad of GI pathologies including inflammation, gastric and intestinal metaplasia, loss of antral identity and perturbations in EEC function. This study represents the utility of human organoids in the identification and modelling of complex pathologies which in turn can alter patient diagnoses and clinical care.

## Figure Legends

**Supplemental Figure 1: (A)** Schematic demonstrating regions of the GI tract and corresponding transcription factors, epithelial markers and hormones.

**Supplemental Figure 2: (A)** Loss of PDX1 results in loss of endocrine cells. Immunofluorescence of Control and PDX1^188delC/188delC^ A staining for secretory protein CHGA, transcription factors PDX1 and NKX2.2 and hormones INS, GLUC, SST. **(B)** Loss of PDX1 results in intestinal or gastric metaplasia in the stomach and intestine, respectively. **(B)** Immunofluorescence and **(C)** Quantification of control, PDX1^188delC/188delC^, and PDX1^188delC/188delC^ CRISPR Corrected antral and post-transplantation intestinal organoids and antral and duodenal biopsies from control, PDX1^188delC/188delC^ A and PDX1^188delC/188delC^ B biopsies stained for epithelial markers CLDN18, CDH1 and transcription factor CDX2. **(D)** Immunofluorescence and **(E)** Quantification of and antral and duodenal biopsies from control, PDX1^188delC/188delC^ A and PDX1^188delC/188delC^ B biopsies stained for intestinal mucin MUC2, gastric mucin MUC5AC and epithelial marker CDH1. All sections counterstained with nuclear DAPI. Scale bars represent 100 μm.

**Supplementary Table 1:** Primary antibody list.

**Supplementary Table 2:** qPCR Primer list.

