## Supplemental Figures for "Using Human Induced Pluripotent Stem Cell Derived Organoids to Identify New Pathologies in Patients with PDX1 Mutations"

### Slide 1
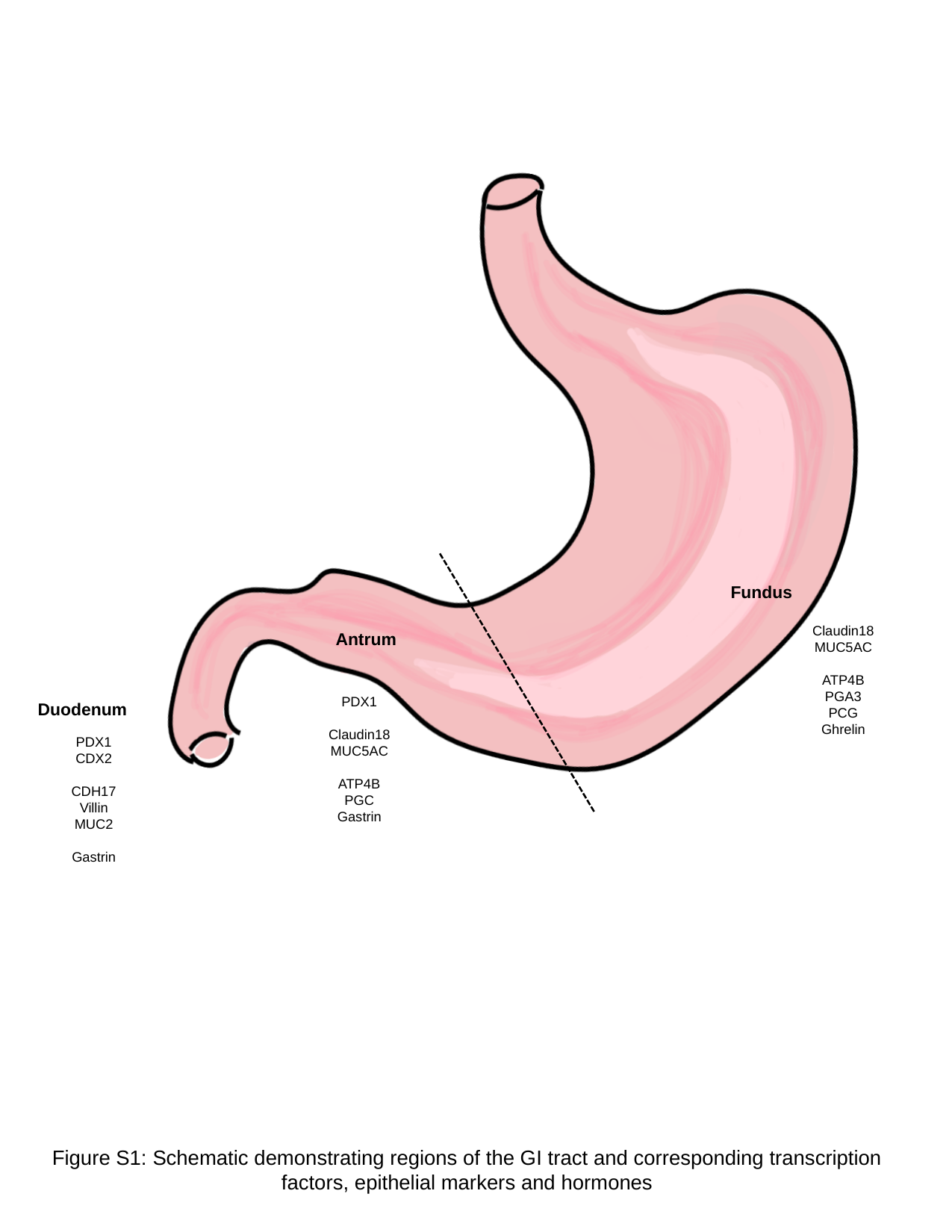

Fundus
Claudin18
MUC5AC
ATP4B
PGA3
PCG
Ghrelin
Antrum
PDX1
Claudin18
MUC5AC
ATP4B
PGC
Gastrin
Duodenum
PDX1
CDX2
CDH17
Villin
MUC2
Gastrin
Figure S1: Schematic demonstrating regions of the GI tract and corresponding transcription factors, epithelial markers and hormones

### Slide 2
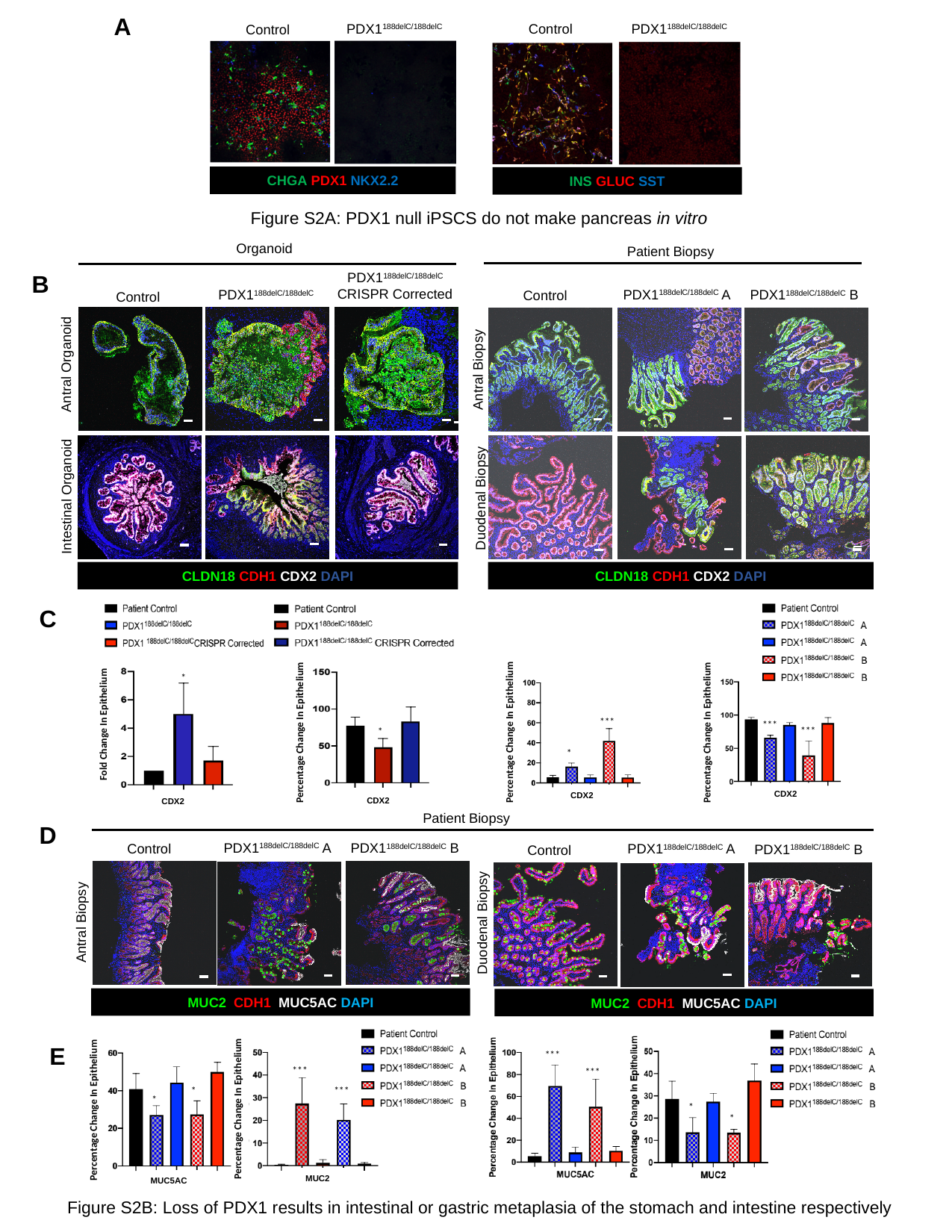

A
CHGA PDX1 NKX2.2
INS GLUC SST
PDX1^188delC/188delC^
PDX1^188delC/188delC^
Control
Control
Figure S2A: PDX1 null iPSCS do not make pancreas in vitro
Organoid
Patient Biopsy
B
Antral Organoid
Antral Biopsy
CLDN18 CDH1 CDX2 DAPI
Intestinal Organoid
Duodenal Biopsy
CLDN18 CDH1 CDX2 DAPI
PDX1^188delC/188delC^
CRISPR Corrected
PDX1^188delC/188delC^ A
PDX1^188delC/188delC^ B
PDX1^188delC/188delC^
Control
Control
*
Percentage Change In Epithelium
CDX2
***
***
Percentage Change In Epithelium
CDX2
*
CDX2
***
*
Fold Change In Epithelium
Percentage Change In Epithelium
CDX2
CDX2
C
Patient Biopsy
D
PDX1^188delC/188delC^ A
PDX1^188delC/188delC^ B
Control
PDX1^188delC/188delC^ A
PDX1^188delC/188delC^ B
Control
Duodenal Biopsy
 Antral Biopsy
MUC2 CDH1 MUC5AC DAPI
MUC2 CDH1 MUC5AC DAPI
 ***
 ***
Percentage Change In Epithelium
MUC2
 *
 *
Percentage Change In Epithelium
MUC5AC
 *
 *
 ***
 ***
E
Figure S2B: Loss of PDX1 results in intestinal or gastric metaplasia of the stomach and intestine respectively

### Slide 3
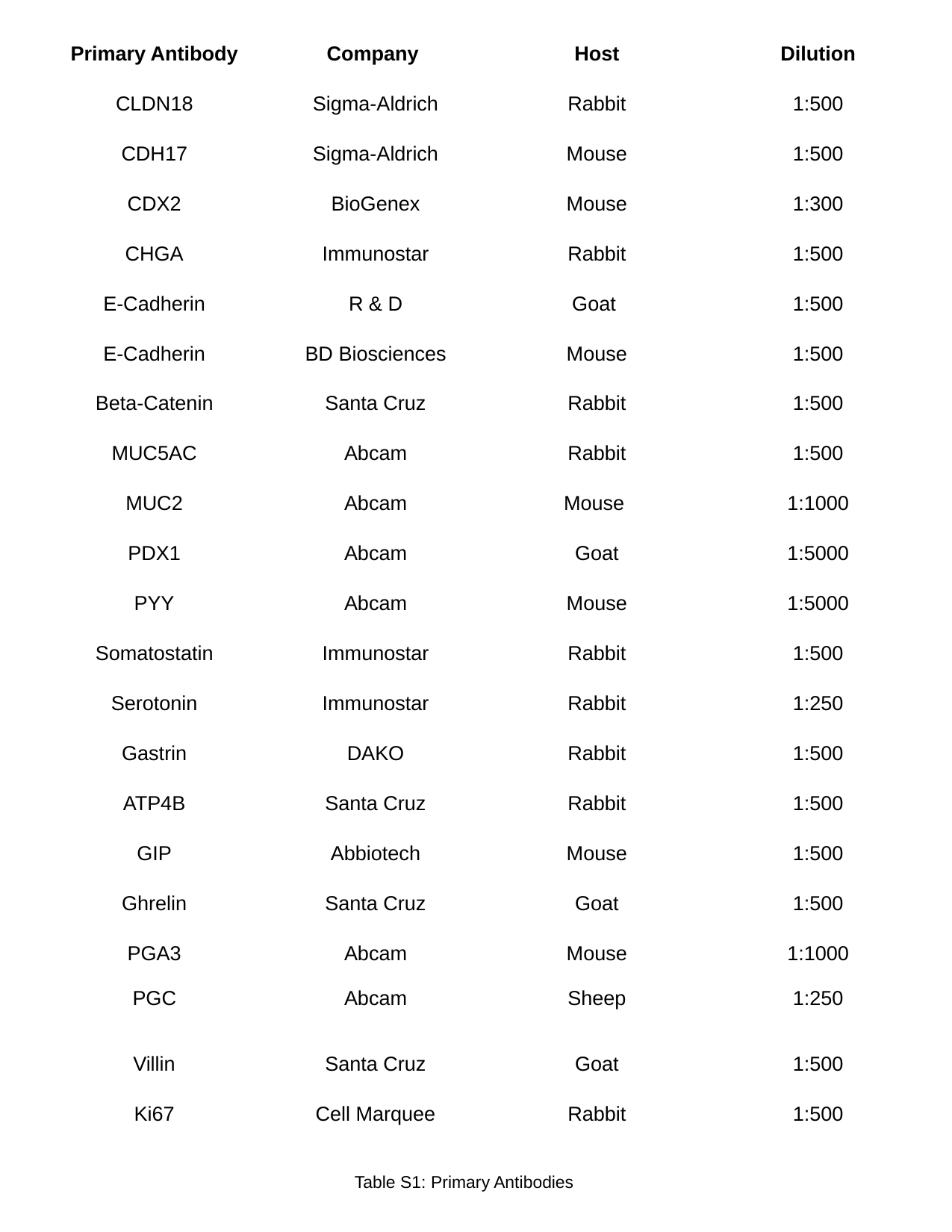

| Primary Antibody | Company | Host | Dilution |
| --- | --- | --- | --- |
| CLDN18 | Sigma-Aldrich | Rabbit | 1:500 |
| CDH17 | Sigma-Aldrich | Mouse | 1:500 |
| CDX2 | BioGenex | Mouse | 1:300 |
| CHGA | Immunostar | Rabbit | 1:500 |
| E-Cadherin | R & D | Goat | 1:500 |
| E-Cadherin | BD Biosciences | Mouse | 1:500 |
| Beta-Catenin | Santa Cruz | Rabbit | 1:500 |
| MUC5AC | Abcam | Rabbit | 1:500 |
| MUC2 | Abcam | Mouse | 1:1000 |
| PDX1 | Abcam | Goat | 1:5000 |
| PYY | Abcam | Mouse | 1:5000 |
| Somatostatin | Immunostar | Rabbit | 1:500 |
| Serotonin | Immunostar | Rabbit | 1:250 |
| Gastrin | DAKO | Rabbit | 1:500 |
| ATP4B | Santa Cruz | Rabbit | 1:500 |
| GIP | Abbiotech | Mouse | 1:500 |
| Ghrelin | Santa Cruz | Goat | 1:500 |
| PGA3 PGC | Abcam Abcam | Mouse Sheep | 1:1000 1:250 |
| Villin | Santa Cruz | Goat | 1:500 |
| Ki67 | Cell Marquee | Rabbit | 1:500 |
Table S1: Primary Antibodies

### Slide 4
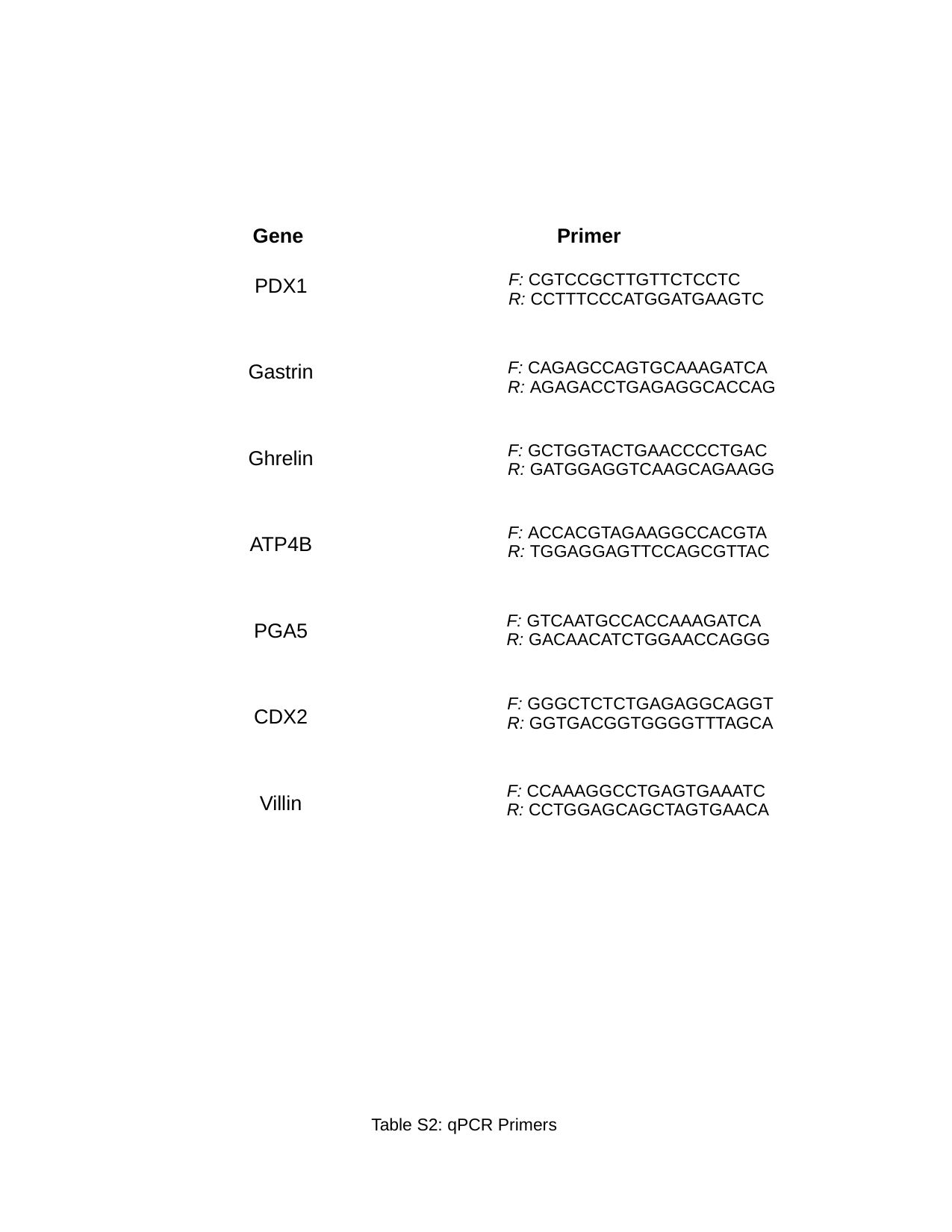

| Gene | Primer |
| --- | --- |
| PDX1 | |
| Gastrin | |
| Ghrelin | |
| ATP4B | |
| PGA5 | |
| CDX2 | |
| Villin | |
| F: CGTCCGCTTGTTCTCCTC R: CCTTTCCCATGGATGAAGTC |
| --- |
| F: CAGAGCCAGTGCAAAGATCA R: AGAGACCTGAGAGGCACCAG |
| --- |
| F: GCTGGTACTGAACCCCTGAC R: GATGGAGGTCAAGCAGAAGG |
| --- |
| F: ACCACGTAGAAGGCCACGTA R: TGGAGGAGTTCCAGCGTTAC |
| --- |
| F: GTCAATGCCACCAAAGATCA R: GACAACATCTGGAACCAGGG |
| --- |
| F: GGGCTCTCTGAGAGGCAGGT R: GGTGACGGTGGGGTTTAGCA |
| --- |
| F: CCAAAGGCCTGAGTGAAATC R: CCTGGAGCAGCTAGTGAACA |
| --- |
Table S2: qPCR Primers
